# ProtAlign-ARG: Antibiotic Resistance Gene Characterization Integrating Protein Language Models and Alignment-Based Scoring

**DOI:** 10.1101/2024.03.20.585944

**Authors:** Shafayat Ahmed, Muhit Islam Emon, Nazifa Ahmed Moumi, Lifu Huang, Dawei Zhou, Peter Vikesland, Amy Pruden, Liqing Zhang

## Abstract

The evolution and spread of antibiotic resistance pose a global health challenge. Whole genome and metagenomic sequencing pose a promising approach to monitoring the spread, but typical alignment-based approaches for antibiotic resistance gene (ARG) detection are inherently limited in the ability to detect new variants. Large protein language models could present a powerful alternative but are limited by databases available for training. Here we introduce ProtAlign-ARG, a novel hybrid model combining a pre-trained protein language model and an alignment scoring-based model to expand the capacity for ARG detection from DNA sequencing data. ProtAlign-ARG learns from vast unannotated protein sequences, utilizing raw protein language model embeddings to improve the accuracy of ARG classification. In instances where the model lacks confidence, ProtAlign-ARG employs an alignment-based scoring method, incorporating bit scores and e-values to classify ARGs according to their corresponding classes of antibiotics. ProtAlign-ARG demonstrated remarkable accuracy in identifying and classifying ARGs, particularly excelling in recall compared to existing ARG identification and classification tools. We also extended ProtAlign-ARG to predict the functionality and mobility of ARGs, highlighting the model’s robustness in various predictive tasks. A comprehensive comparison of ProtAlign-ARG with both the alignment-based scoring model and the pre-trained protein language model demonstrated the superior performance of ProtAlign-ARG.

## Introduction

Antibiotic resistance is a major threat to public health. Every year, antibiotic-resistant human pathogens claim the lives of an estimated 700,000 people worldwide, with this number expected to rise to 10 million per year by 2050^1,2^. Antibiotic-resistance genes (ARGs), i.e., genes carried by bacteria that confer resistance to antibiotics, can be shared between bacteria and then further transmitted among animals, humans, and various environments^3–7^. Thus, there have been calls for the advancement of approaches for monitoring the emergence and movement of ARGs across One Health sectors^8^.

With the advent of next-generation DNA sequencing technologies, databases of bacterial genomes and metagenomes originating from various One Health sources continue to grow. This presents the opportunity to tap into such data to advance ARG monitoring. Traditionally, ARGs are identified from DNA sequencing data via alignment to existing ARG databases, such as the Comprehensive Antibiotic Resistance Database (CARD)^9^. Sequence alignment entails the computational arrangement of nucleotide or amino acid sequences to determine regions of similarity, which may indicate functional, structural, or evolutionary relationships among the sequences. Unfortunately, alignment-based approaches are inherently limited, particularly in their reliance on existing databases and their inability to capture remote homologs, i.e., instances where evolutionary relationships between genes or proteins have significantly diverged over time^10^. Further, alignment-based methods are highly sensitive to the selection of similarity thresholds, resulting in false negatives if too stringent or false positives if too liberal. This ambiguity poses challenges for users, but also inherently limits the ability to detect novel ARGs.

Recognizing these constraints underscores the importance of exploring alternative methodologies to enhance the accuracy and precision of ARG prediction for antibiotic resistance monitoring. Despite the broad adoption of alignment-based algorithms in ARG classification, it is clear that this approach alone doesn’t provide comprehensive insights for accurate ARG class classification. Moreover, aligning against large databases is time-consuming^11^, often requiring hours to days to align terabytes of sequence data to databases of several gigabytes in size^12–14^.

Deep learning-based tools have the potential to greatly advance One Health monitoring of ARGs, presenting a means to improve the efficiency and accuracy of ARG detection and classification from DNA sequencing data, while also enabling the detection of new/emergent ARGs that are not yet described in publicly available databases. Deep learning, particularly with protein language model embeddings, offers a more nuanced representation of protein sequences, excelling in contextualizing these sequences and uncovering complex patterns missed by conventional methods^15^. Recently, deep learning-based tools have demonstrated promise for ARG identification and classification. Deep-ARG^16^ is one such tool that uses deep learning and considers a dissimilarity matrix for ARG identification. HMD-ARG^17^ was developed more recently and offers a hierarchical multi-task classification model using a convolutional neural network(CNN). ARG-SHINE^18^ utilizes a machine learning approach to ensemble three component methods for predicting ARG classes. However, protein language models have only begun to be tested for ARG detection. Recently, a transformer-based protein language model demonstrated strong generalization capabilities, meaning it could accurately learn patterns from a limited number of training sequences and apply this knowledge to effectively identify and classify new, unseen protein sequences^19^. Transformer models use a self-attention mechanism to capture long-range dependencies and contextual relationships, enhancing their performance on sequence data.

The abundance of protein sequences translated from next-generation DNA sequencing data has revealed complex molecular relationships and intricate interactions between amino acids far apart in the protein’s three-dimensional structure. Protein sequences also inherently encode crucial structural features, including secondary structures and motifs, which guide protein folding and function^20^. Protein language models provide a powerful approach to deciphering such complexity. In particular, these models excel in capturing intricate patterns and motifs across diverse gene types, providing a systematic way to understand the nuanced language of protein sequences. Protein language models, trained on millions of protein sequences, have demonstrated promise for developing ARG prediction models. Our preliminary study demonstrated the efficacy of pre-trained protein language models for ARG identification and classification tasks^21^. Recently PLM-ARG^22^ has also proven capable of using protein language models to predict ARGs. We found that integrating language models and transfer learning, a technique where a model developed for a specific task is reused as the starting point for a model on a second task^23^, could be a promising approach to ARG identification and classification while mitigating false negatives. However, a notable challenge remains: deep learning models exhibit suboptimal performance when confronted with limited training data, which hampers their ability to support generalized predictions. In such cases of insufficient training data, alignment-based scoring can outperform pre-trained protein language model (PPLM) based prediction.

Here we introduce ProtAlign-ARG as a novel model for identifying and classifying ARGs in protein sequence data that leverages the strengths of PPLM-based prediction with alignment-based scoring. This hybrid solution aims to enhance predictive accuracy, particularly in scenarios with limited training samples and diverse datasets. To demonstrate the ProtAlign-ARG pipeline, we carried out a comprehensive comparison with existing pipelines, validating the superiority of our proposed model in overcoming challenges posed by inadequate training data. We further demonstrate the capability of ProtAlign-ARG to characterize ARG mobility, including differentiating intrinsic ARGs, along with improvements in ARG classification according to antibiotic resistance class.

## Model Development

### Data Curation

For ARG data, we utilized HMD-ARG-DB^17^ as it is one of the largest repositories of ARGs and one that captures the most comprehensive annotations across various dimensions. HMD-ARG-DB was curated from seven widely-used databases, namely AMRFinder^24^, CARD^25^, ResFinder^26^, Resfams^27^, DeepARG^16^, MEGARes^28^, and Antibiotic Resistance Gene-ANNOTation^29^, containing over 17,000 ARG sequences distributed among 33 antibiotic-resistance classes. As 19 classes in HMD-ARG-DB have only a few genes in their groups, we focused on the 14 most prevalent classes for developing the prediction models.

To curate the non-ARG dataset, we downloaded the entire Uniprot dataset while excluding sequences labeled as ARGs and performed diamond alignment with the HMD-ARG-DB. Sequences with an e-value greater than 1e-3 and a percentage identity below 40% were classified as non-ARGs. This enabled focus on non-ARG sequences demonstrating some level of similarity to ARG sequences. The goal was to enhance the model’s capability to identify ARGs, even in scenarios where such similarities exist^17^. Furthermore, it facilitated comparisons with other state-of-the-art tools such as DeepARG^16^ and HMD-ARG^17^.

For comparing our performance with other ARG annotation and classification tools we additionally performed experiments using the COALA (COllection of ALl Antibiotic resistance gene databases) dataset^18^. The dataset was collected from 15 published related databases including CARD^9^, ResFinder^26^, ResfinderFG^30^, ARDB^31^, MEGARes^28^, NDARO^32^, ARG-ANNOT^29^, Mustard^33^, FARME^34^, SARG(v2)^35^, Lahey list of beta-lactamases^36^, BLDB^37^, LacED^38,39^, CBMAR^40^, MUBII-TB-DB^41^, and u-CARE^42^. Hamid et al.^43^ collected protein sequences from these databases with their annotations. The dataset has sixteen drug resistance classes and in total 17023 ARG sequences.

### Data Partition

We partitioned the dataset into training, validation, and testing sets based on a specific similarity threshold. This threshold represented the maximum allowed similarity between sequences in the two sets, ensuring that the training and testing data were distinct enough to prevent biased accuracy metrics, and to assess the accuracy of each model on unseen data. Although CDHIT^44^ and MMseq^45^ are commonly used for data partitioning, they cannot guarantee that training and testing data have the specified maximum similarity. In contrast, GraphPart^46^, a recently developed tool, has proven to be highly effective in precise separation and retain most sequences until reaching the desired similarity threshold. Here we used both CDHIT and GraphPart for data partitioning and clustering. Using CDHIT^44^ with a 40% threshold similarity on the HMD-ARG-DB dataset, we obtained 721 clusters that were partitioned into training and testing data with an 80:20 ratio. However, as shown in Figure 5, a comparison of training and testing data revealed that more than 50% of sequences between the training and testing sets partitioned by CDHIT had similarity over 40% (200 were > 90%). In contrast, GraphPart provided exceptional partitioning precision and is therefore used for partitioning 80% of the sequences for training and validation and 20% for the testing. Table 1 shows the distribution of ARG counts in training and testing sets using Graphpart for 40% and 90% similarity thresholds.

**Table 1.**
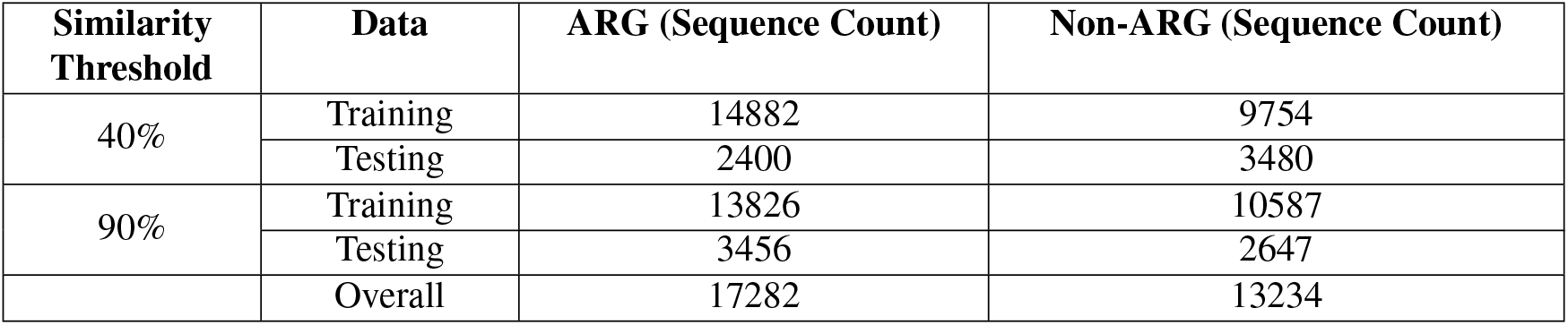
Dataset split for HMD-ARG-DB^17^ using Graphpart with 40% and 90% threshold.

### ProtAlign-ARG tasks and models

ProtAlign-ARG comprises four distinct models, each dedicated to a specific task, including 1) ARG Identification, 2) ARG Class Classification, 3) ARG Mobility Identification, and 4) ARG Resistance Mechanism. Figure 1 presents an overview of the architecture of how these four models were developed, highlighting their respective designs and interrelationships. ProtAlign-ARG utilizes both pre-trained protein language models and alignment-based scoring model to ensure robust performance in these tasks. Detailed descriptions of these components are provided in the following sections.

**Figure 1.**
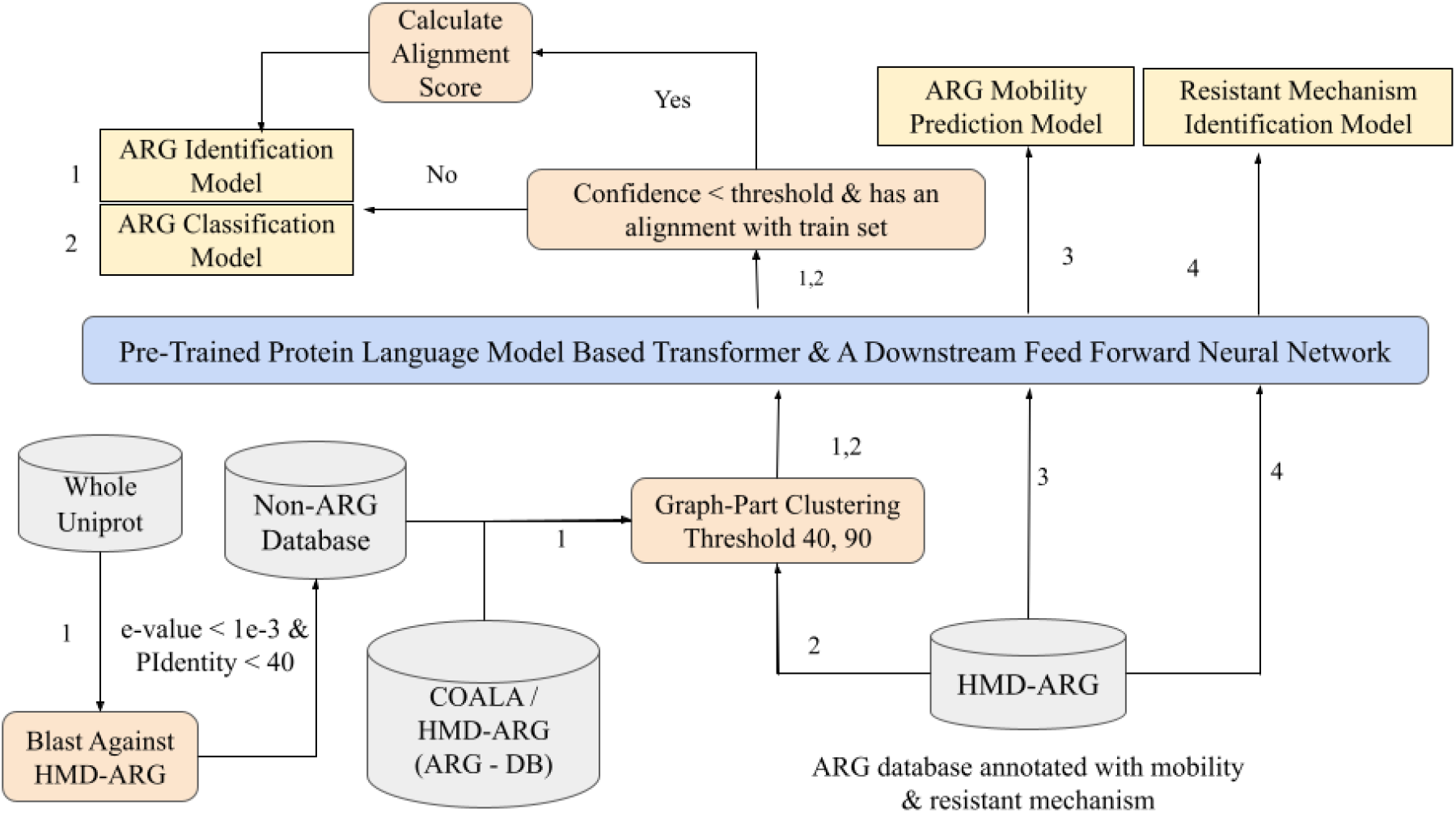
The pipeline for ARG Identification & Classification. Each number (1-4) corresponds to the four distinct models incorporated in the pipeline.

### Leverage pre-trained language Model for ARG identification and classification

At the time of our work, many protein language models are available, such as ProtXL, ProtElectra, and ProtT5. We selected the ProtAlbert model due to its superior performance. Notably, ProtAlbert outperformed its counterparts while maintaining a relatively modest size, comprising 224 million parameters. This choice strategically balanced between optimal performance and computational efficiency, especially when compared to larger models that are more time-consuming. ProtAlbert, with its 12-layer architecture, undergoes training on UniRef100. For our downstream prediction task, we harnessed the pre-trained ProtAlbert model following the official GitHub repository provided by ProtTrans^20^. The training unfolds in two stages: an initial phase on sequences of up to 512 characters for 150,000 steps, succeeded by another 150,000 steps on longer sequences, spanning up to 2000 characters.

The pre-trained ProtAlbert model imparts its knowledge to the ARG classification task through transfer learning embeddings. ProtAlbert offers two methods: per-residue, which treats each amino acid separately, and per-protein, which views the entire protein sequence as one unit. For ARG identification and classification, we chose the per-protein method. This approach recognizes proteins as integrated units with specific structures and functions. It simplifies the representation, making it less computationally complex and better at handling sequence variations. The per-residue method gives detailed amino acid information but can be more complex and sensitive to sequence changes. Thus, the per-protein method balanced biological relevance with computational efficiency. As demonstrated in Figure 2, we extract embeddings from the model’s final layer and consolidate them into a fixed-size representation for all proteins. ProtAlbert employs mean pooling for this purpose. The resultant vector serves as the input for a single feed-forward layer in the ARG identification or classification task, featuring 32 neurons.

**Figure 2.**
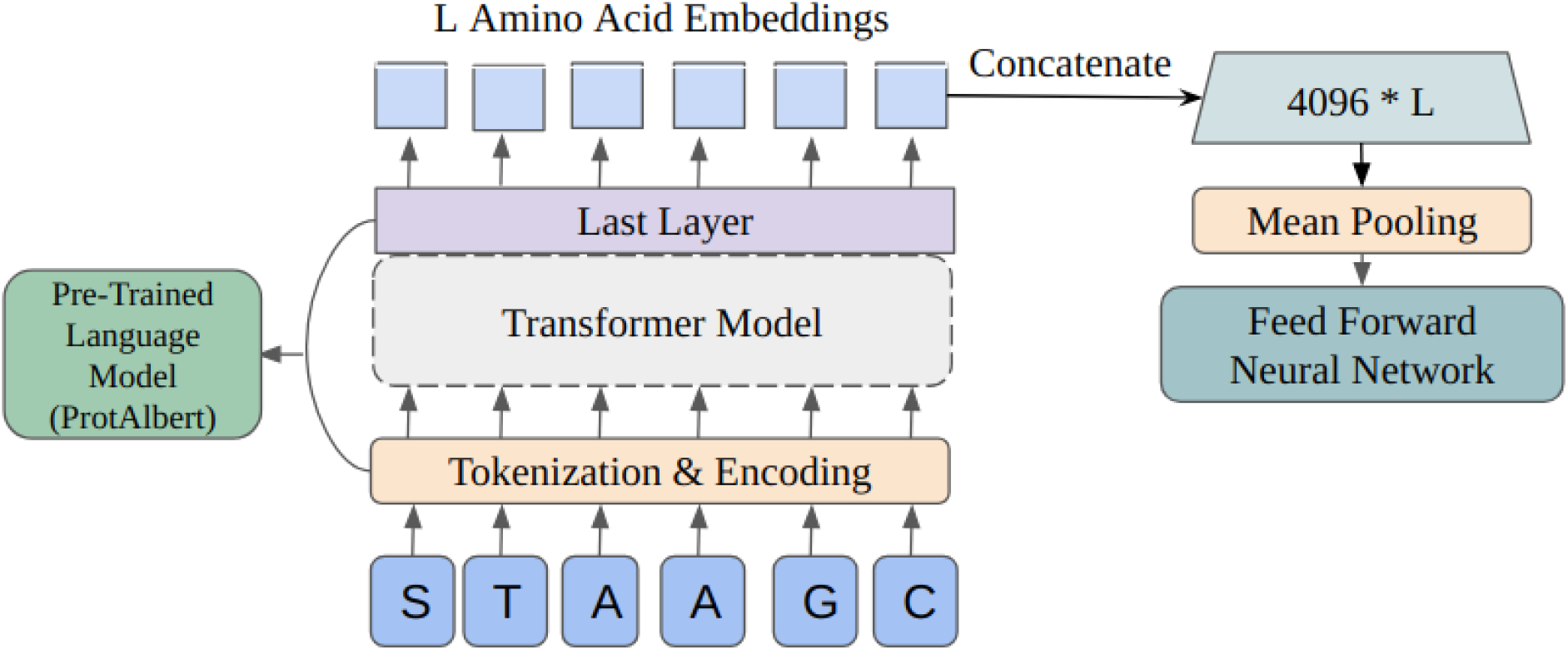
Pre-trained protein language model for ARG Identification & Classification.

### Alignment-based scoring model

Deep learning models are effective in learning from large data sets but may not perform well with small amounts of data. In contrast, alignment-based models focus on sequence similarities and can perform well even if there is a limited number of sequences. So while deep learning performs well on large amounts of sequence data, alignment-based models are often better for smaller data sets. Here we combine both models for ARG identification and prediction, leveraging the advantages of both models. For the alignment-based scoring, we utilized DIAMOND^47^ for matching, following a modified version of the ARG-KNN model proposed by the paper ARG-SHINE^18^.

Initially, we align the query sequence with the training data using DIAMOND, setting an e-value threshold of *<* 1*e*− 3 to identify similar sequences (homologs). If a query sequence fails to align with any sequence in the training dataset, we cannot employ the alignment-based scoring to label it. We switch to PPLM prediction for those sequences. We applied this scoring method to both ARG identification and classification.

For each query sequence, we computed alignment scores for each label. The label with the highest score is assigned to the query sequence. For a given query sequence, denoted as *p*_*q*_, where *q* is the number of sequences from the same label. The score for the label *C*_*i*_, *S*(*C*_*i*_, *p*_*q*_) is defined by the following equation:

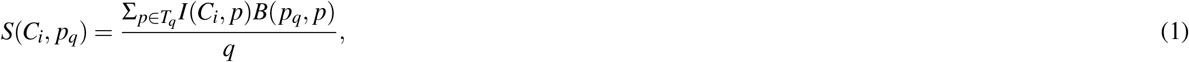

where *T*_*q*_ represents the set of proteins associated with label *C*_*i*_ and their bit scores for *p*_*q*_, *p* signifies any protein in *T*_*q*_, *I*(*C*_*i*_, *p*) is a binary indicator indicating whether *p* belongs to the label *C*_*i*_, and *B*(*p*_*q*_, *p*) signifies the bit score of the alignment between protein *p*_*q*_ and *p*. The label with the highest score is considered the result of the similarity model.

### ProtAlign-ARG

ProtAlign-ARG synergizes PPLM-based scoring with alignment-based scoring to enhance the accuracy of ARG identification and classification. Deep learning models, particularly PPLMs, are known to excel in data-rich environments; however, they tend to struggle with accuracy when the available data are sparse or when the model’s predictive confidence is low. In our methodology, we introduce confidence thresholds as a metric for evaluating the reliability of PPLM-based predictions. These thresholds—95%, 90%, 80%, 70%, 60%, 50%, 40%, 30%, and 20%—represent the model’s certainty in its classification. We assessed the accuracy of predictions falling below each threshold and observed a marked decline in reliability (Figure 3). Specifically, predictions with less than 90% confidence yielded a 45% accuracy rate, indicating that over half of these low-confidence predictions were incorrect. Consequently, we established 90% as the minimum confidence threshold for utilizing PPLM-based classification for both ARG identification and class classification. We defaulted to alignment-based scoring for predictions falling below this 90% confidence threshold. This scoring system is more robust in scenarios with limited data as it does not rely on the voluminous training datasets that PPLM requires. It improves prediction reliability by comparing the query sequence against a database of known sequences.

**Figure 3.**
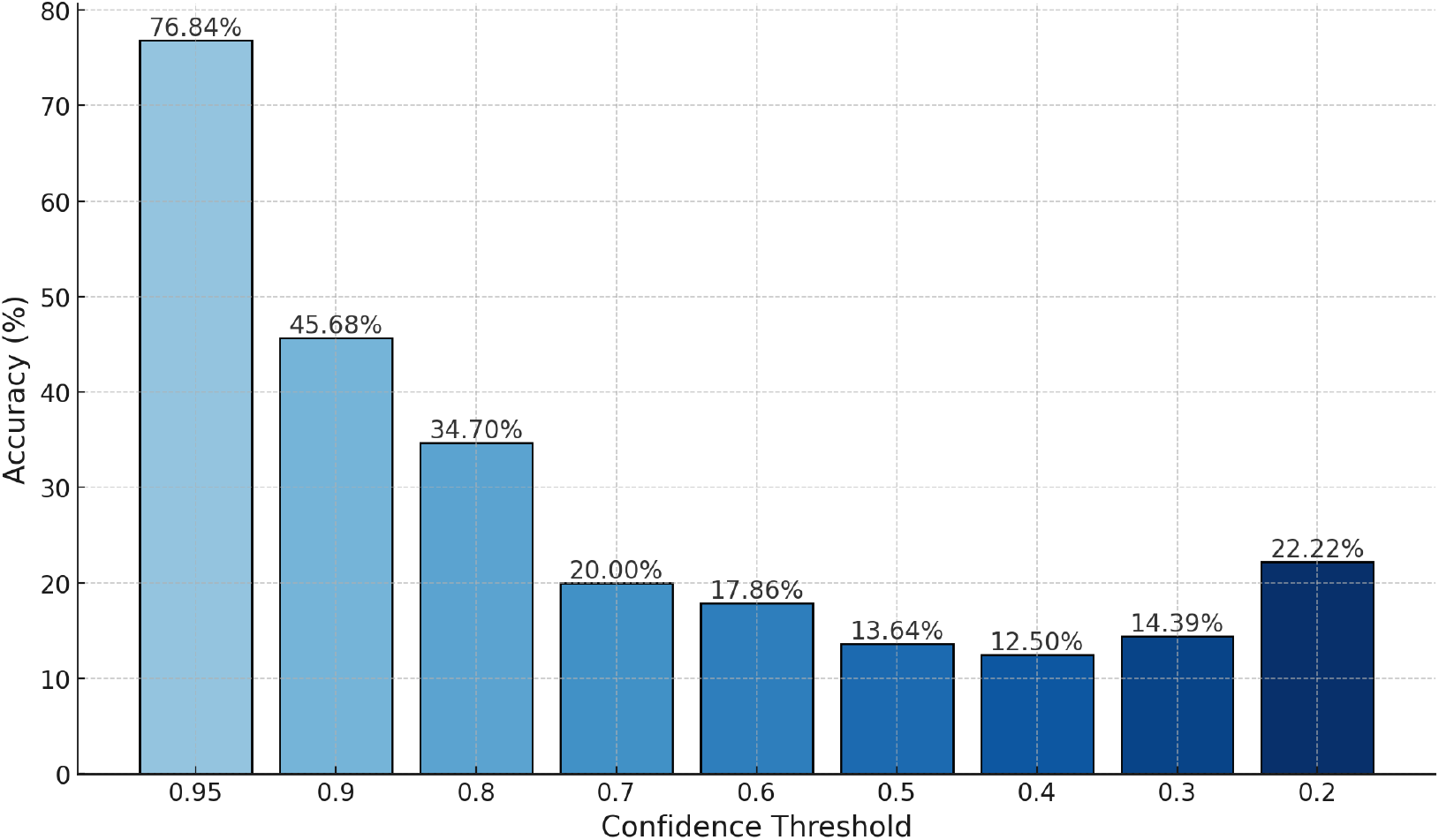
Percentage of accuracy for predictions below the indicated confidence thresholds for PPLM-based ARG class classification.

When the query sequence does not find a match within the training data, which is a situation expected to arise with novel or rare ARGs, PPLM is then applied for the final classification. The PPLM, leveraging its comprehensive understanding of protein sequence language, is a decision-making tool in such cases. Figure 3 illustrates the relationship between the various confidence thresholds and the corresponding accuracy of predictions.

### Code Availability

The complete code, data, and, results are included in the repository - https://github.com/Shafayat115/ProtAlign-ARG.

## Results & Discussions

### ARG Identification Models

We conducted training and testing using two different setups, employing both a 40% threshold and a 90% threshold to assess the performance of ProtAlign-ARG. Table 2 provides an overview of the accuracy achieved in each setup, distinguishing between ProtAlign-ARG, PPLM, and alignment-scoring-based prediction. The ARG identification model of ProtAlign-ARG consistently outperformed alignment-scoring-based ARG identification models, even when the training and testing sets had a similarity of under 40%. It achieved an overall F1-score of 84%. However, the ARG identification model of ProtAlign-ARG did not substantially improve overall accuracy relative to the PPLM or alignment-scoring-based ARG identification models. This result was as expected, since both of the labels applied in this study, “ARG” and “non-ARG”, had ample data in both the training and test sets for this portion of the study.

**Table 2.** Accuracy for component models on ARG Identification on HMD-ARG-DB^17^ using 40 and 90 percent similarity threshold using GraphPart^46^.

| Train-Test Similarity | Model | Precision | Recall | F1-Score |
| --- | --- | --- | --- | --- |
| 40% | PPLM | 0.85 | 0.83 | **0.84** |
|  | Alignment | 0.55 | 0.55 | 0.55 |
|  | ProtAlignARG | **0.86** | **0.84** | **0.84** |
| 90% | PPLM | 0.94 | **0.98** | **0.97** |
|  | Alignment | 0.71 | 0.71 | 0.71 |
|  | ProtAlignARG | **0.95** | 0.96 | 0.96 |

### Classification of ARGs According to Resistance Class

Table 3 shows that the PPLM excelled in achieving better recall compared to the alignment-based scoring model, but it lagged in terms of precision. ProtAlign-ARG, on the other hand, struck a balance by achieving a high level of precision and recall, leading to an overall improvement in accuracy relative to both alternative models.

**Table 3.** Accuracy for the PPLM, Alignment-scoring-based model, and ProtAlign-ARG for ARG class classification on HMD-ARG-DB at 40% threshold.

| Model | Metric | Precision | Recall | F1-Score |
| --- | --- | --- | --- | --- |
| PPLM | Macro | 0.58 | 0.62 | 0.56 |
|  | Weighted | 0.88 | 0.83 | 0.83 |
| Alignment-Scoring | Macro | **0.69** | 0.52 | 0.58 |
|  | Weighted | **0.95** | 0.76 | 0.84 |
| ProtAlignARG | Macro | 0.63 | **0.67** | **0.64** |
|  | Weighted | 0.90 | **0.89** | **0.89** |

Figure 4 shows that the PPLM performed well for ARGs encoding resistance to Aminoglycoside, Beta-lactam, Fosfomycin, Glycopeptide, Tetracycline, and Trimethoprim, with high scores across all metrics. The alignment-based scoring model exhibited high precision and recall for certain classes, such as Aminoglycoside (precision 1 and recall 0.9) and Trimethoprim (precision 1 and recall 1), but performed poorly for Polymyxin and Rifampin (precision and recall 0 for both). The ARG class prediction model of ProtAlign-ARG performed well for classes like aminoglycoside (F1-score 0.9) and beta_lactam (F1-score 1). It also improved the performance of the PPLM model for Quinolone, Sulfonamide, and MLS class for the F1-score (0.5,0.8, and 0.8 respectively).

**Figure 4.**
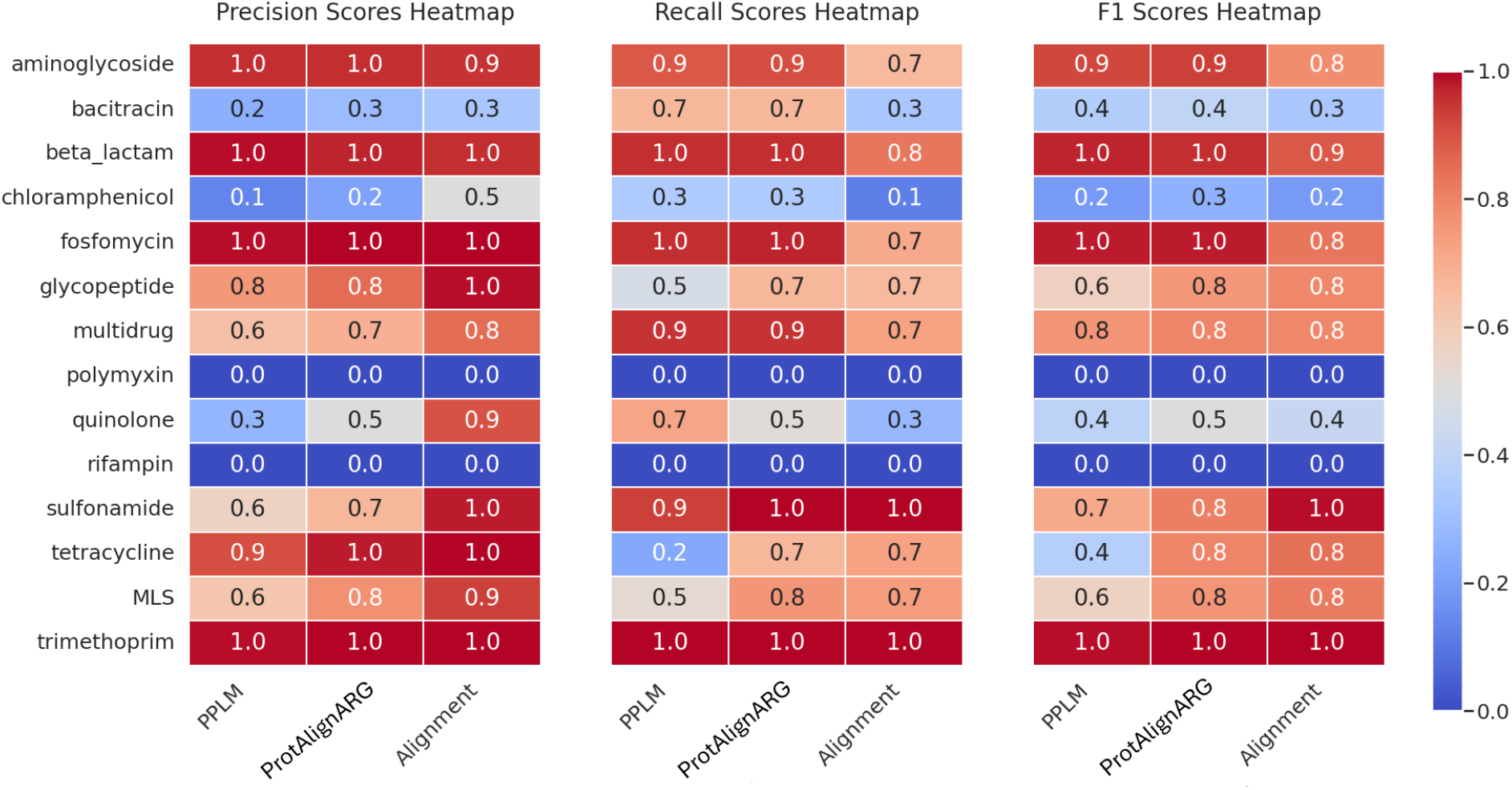
ARG class prediction accuracy for the PPLM, Alignment-scoring-based model, and ProtAlign-ARG based on precision, recall, and F1-score.

**Figure 5.**
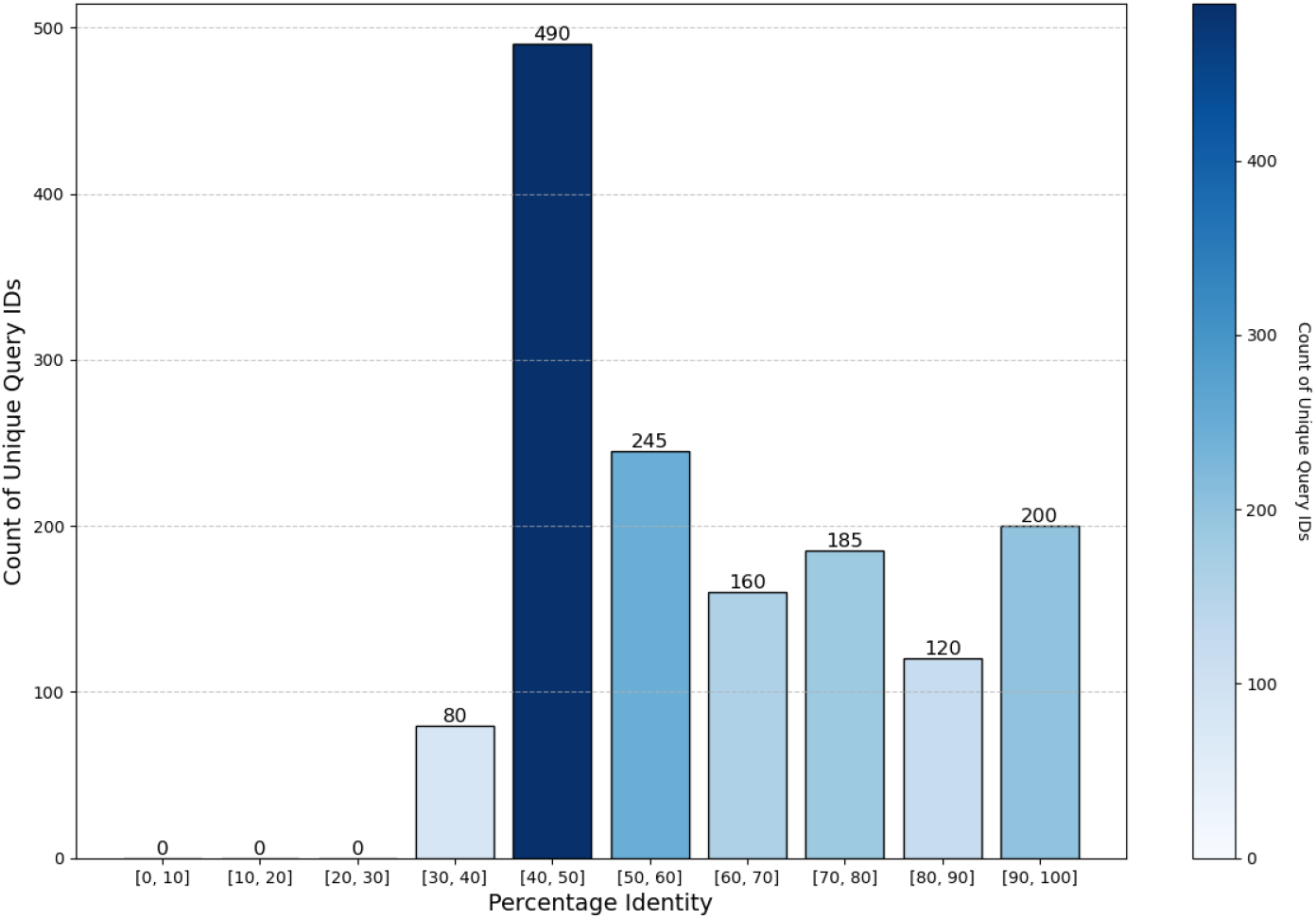
Distribution of sequence count across different similarity thresholds between the training and testing set achieved by CDHIT clustering.

Additionally, we conducted tests using a 90% similarity threshold. This was particularly important because 19 of the ARG class samples had just 219 sequences in total. Table 4 presents the performance of the three models in this scenario. In this case, the alignment-scoring-based model outperformed both the PPLM and the ProtAlign-ARG in precision. The PPLM tends to perform poorly when the sample size is limited (e.g., Kasugamycin and Quinolone antibiotic classes). When we experimented with all 33 ARG classes, including the 19 ARG classes with just only 219 sequences in total, the accuracy dropped significantly for the PPLM model, whereas the alignment-scoring model performed much better. However, when we considered the 90% similarity threshold only for the remainder of the 14 frequent ARG classes, the PPLM outperformed the alignment-scoring-based model. Our hybrid ProtAlign-ARG achieved a 78% F1-score, consistent with the alignment-scoring-based model.

**Table 4.** Accuracy for the different component models for ARG class classification on HMD-ARG-DB at 90% threshold.

| Model | Metric | Precision | Recall | F1-Score |
| --- | --- | --- | --- | --- |
| PPLM | Macro | 0.41 | 0.45 | 0.42 |
|  | Weighted | 0.96 | 0.97 | 0.97 |
| Alignment-Scoring | Macro | 0.80 | 0.80 | 0.78 |
|  | Weighted | 0.98 | 0.98 | **0.98** |
| ProtAlign-ARG | Macro | 0.80 | 0.79 | 0.78 |
|  | Weighted | 0.98 | 0.98 | **0.98** |

**Table 5.** Accuracy for the different component models for ARG class classification on COALA90 dataset.

| Models | Accuracy | Precision | Recall | F1-score |
| --- | --- | --- | --- | --- |
| PPLM | Macro | 0.68 | 0.65 | 0.67 |
|  | Weighted | 0.82 | 0.81 | 0.81 |
| Alignment-Scoring | Macro | 0.79 | 0.70 | 0.71 |
|  | Weighted | 0.89 | 0.74 | 0.80 |
| ProtAlign-ARG | Macro | **0.86** | 0.81 | **0.83** |
|  | Weighted | 0.85 | **0.84** | **0.84** |

Furthermore, for comparing our model’s accuracy with existing state-of-the-art models like DeepARG^16^, ARG-SHINE^18^, and HMD-ARG^17^, we divided the COALA dataset^18^, categorizing it into three groups: “No alignment,” “Less than 50% similarity between training and testing sets,” and “Greater than 50% threshold.” The COALA90 dataset was prepared using the CD-HIT tool, which could not maintain a strict threshold between cluster similarity. In this dataset, the performance of the PPLM deteriorated, especially for the ARG classes streptogramin, rifamycin, and bacitracin, which had just 26, 29, and 127 sequences and achieved an overall F1-score of 0.67. While ProtAlign-ARG outperformed both of them. The model achieved an F1-score of 0.81 even outperforming the alignment-based scoring model that achieved an overall accuracy of F1-score of 0.71.

We also divided the test dataset into three subsections based on the alignment with the training sequences: those with no alignment, those with less than 50% alignment, and those with greater than 50% alignment using the CDHIT tool. Table 6 shows the accuracies achieved by the PPLM and the ProtAlign-ARG for these test sets. We observed that for cases with no alignment and very low alignment, the PPLM performed better than the alignment-scoring-based model and ProtAlign-ARG achieved an overall F1-score of 0.42 and 0.56 respectively outperforming both of them. For alignment over 50%, the ProtAlign-ARG achieved an F1-score of 0.91 whereas the PPLM and the alignment-scoring-based model achieved 0.72 and 0.82 respectively.

**Table 6.** Accuracy for the different component models for ARG class classification on COALA dataset for varying percentage similarity.

| Models | Similarity | Accuracy | Precision (%) | Recall (%) | F1-score (%) |
| --- | --- | --- | --- | --- | --- |
| PPLM | Greater 50 | Macro | 74 | 71 | 72 |
|  |  | Weighted | 90 | 91 | 90 |
|  | Less 50 | Macro | 58 | 53 | 55 |
|  |  | Weighted | 72 | 73 | 72 |
|  | No Alignment | Macro | 42 | 47 | **42** |
|  |  | Weighted | 41 | 45 | **41** |
| Alignment-Scoring | Greater 50 | Macro | 85 | 81 | 82 |
|  |  | Weighted | 93 | 92 | 92 |
|  | Less 50 | Macro | 46 | 41 | 42 |
|  |  | Weighted | 72 | 65 | 68 |
|  | No Alignment | Macro | 0 | 0 | 0 |
|  |  | Weighted | 0 | 0 | 0 |
| ProtAlign-ARG | Greater 50 | Macro | 94 | 89 | **91** |
|  |  | Weighted | 95 | 95 | **95** |
|  | Less 50 | Macro | 56 | 56 | **56** |
|  |  | Weighted | 73 | 73 | **73** |
|  | No Alignment | Macro | 42 | 47 | **42** |
|  |  | Weighted | 41 | 45 | **41** |

### Comparison with Other Methods

#### ARG Identification

In our comprehensive comparison with other state-of-the-art ARG identification tools, ProtAlign-ARG demonstrated superior performance across all metrics when conducted using the HMD-ARG-DB. To ensure the robustness and reliability of our results, we employed a rigorous 5-fold cross-validation approach^48^. This method involves partitioning the dataset into five distinct subsets, wherein each fold, one subset is used for testing and the remaining four for training. This process is repeated five times, with each subset used exactly once as the test set. Furthermore, to account for variability and ensure statistical significance, we repeated the entire cross-validation procedure 10 times. This repetition helps in averaging out any anomalies or fluctuations in the data, providing a more accurate estimate of the model’s performance. Our extensive testing revealed that ProtAlign-ARG consistently achieved an impressive accuracy of 97%, as detailed in Table 7. This high level of accuracy indicates the model’s exceptional capability to correctly identify ARGs, even in complex and diverse datasets. Compared to other leading ARG identification tools, ProtAlign-ARG’s performance not only highlights its precision but also its robustness and generalizability across different subsets of data. The balanced F1 score showcases the model’s overall reliability in ARG identification tasks.

**Table 7.** Accuracy for the different component models for ARG Identification on HMD-ARG-DB.

| Models | Precision | Recall | F1-Score |
| --- | --- | --- | --- |
| HMD-ARG | 0.939 | 0.971 | 0.948 |
| CARD | 0.999 | 0.421 | 0.592 |
| DeepARG | 0.998 | 0.93 | 0.963 |
| AMRPlusPlusb | 0.867 | 0.449 | 0.592 |
| Meta-MARC | 0.847 | 0.85 | 0.848 |
| ProtAlign-ARG | 0.97 | 0.98 | 0.97 |

#### ARG Classification

When we compared ProtAlign-ARG’s ARG class classification model on the COALA dataset, it outperformed most alignment-based classification tools and delivered comparable performance to deep learning-based tools. We evaluated using the COALA90 dataset, which was generated using CDHIT clustering. Although the model (ARG class classification model of ProtAlign-ARG) was trained on the same set of training data, however, the test set was three times larger than that used by other models. Table 8, summarizes the performance of our model relative to other models. We can see that ProtAlign-ARG achieved an F1-score of 0.83.

**Table 8.** Accuracy for the different models for ARG Classification on COALA90 dataset.

| Models | Macro Avg. Score | Weighted Avg. Score |
| --- | --- | --- |
| BLAST best hit | 0.8258 | 0.8423 |
| DIAMOND best hit | 0.8103 | 0.8423 |
| DeepARG | 0.7303 | 0.8419 |
| HMMER | 0.4499 | 0.4916 |
| TRAC | 0.7399 | 0.8097 |
| ARG-SHINE | 0.8555 | 0.8591 |
| PPLM Model | 0.67 | 0.81 |
| Alignment-Score | 0.71 | 0.80 |
| ProtAlign-ARG | 0.83 | 0.84 |

We furthermore compared ProtAlign-ARG using the three splits of the COALA90 dataset. In this case, we reported the overall accuracy of the ARG class classification model of ProtAlign-ARG, emphasizing that this model was tested on test sets three times larger than other tools (Table 9). ProtAlign-ARG achieved an overall F1-score of 0.45, 0.73, and 0.95 respectively for the no-alignment, alignment less and greater than 50 percentage splits.

**Table 9.** Accuracy for the different component models for ARG Classification on three splits of COALA90 dataset.

| Models | No-Alignment | Alignment <50 | Alignment >50 |
| --- | --- | --- | --- |
| BLAST | 0 | 0.6243 | 0.9542 |
| DIAMOND | 0 | 0.5740 | 0.9534 |
| DeepARG | 0 | 0.5266 | 0.9419 |
| HMMER | 0.0563 | 0.2751 | 0.6051 |
| TRAC | 0.3521 | 0.6124 | 0.9199 |
| ARG-SHINE | 0.4648 | 0.6864 | 0.9558 |
| PPLM Model | 0.45 | 0.73 | 0.91 |
| Alignment | 0 | 0.65 | 0.92 |
| ProtAlignARG | 0.45 | 0.73 | 0.95 |

We also show the accuracy of the ARG class classification model of ProtAlign-ARG using a 5-fold cross-validation approach on the HMD-ARG-DB in Table 10. ProtAlign-ARG consistently outperformed the other models in this evaluation. The model hit an average F1-score of 0.96 whereas the second-best-performing model HMD-ARG got an overall F1-score of 0.89.

**Table 10.** Accuracy for the state-of-the-art tools and ARG class classification model of ProtAlign-ARG on HMD-ARGDB using 5-fold cross validation.

| Models | Precision | Recall | F1-Score |
| --- | --- | --- | --- |
| HMD-ARG | 0.950 | 0.847 | 0.893 |
| DeepARG | 0.914 | 0.757 | 0.814 |
| CARD | 0.983 | 0.452 | 0.585 |
| META-MARC | 0.750 | 0.782 | 0.745 |
| ProtAlign-ARG | 0.97 | 0.94 | 0.96 |

Additionally, we expanded the scope of our prediction tasks to include ARG resistance mechanism prediction and mobility identification. In both cases, PPLM achieved better performance than other state-of-the-art tools when using 5-fold cross-validation (Table 12 & 11).

**Table 11.** Accuracy for resistance mechanism classification on HMD-ARGDB using 5-fold cross-validation for different models.

| Models | Precision | Recall | F1-Score |
| --- | --- | --- | --- |
| HMD-ARG | 0.867 | 0.768 | 0.795 |
| CARD | 0.832 | 0.476 | 0.566 |
| PPLM | 0.97 | 0.96 | 0.97 |

**Table 12.** Accuracy for mobility identification on HMD-ARGDB using 5-fold cross validation.

| Models | Precision | Recall | F1-Score |
| --- | --- | --- | --- |
| HMD-ARG | 0.964 | 0.89 | 0.926 |
| PPLM | 0.98 | 0.99 | 0.99 |

For the resistance mechanism classification task, PPLM reached an F1-score of 0.97 whereas the closest one HMD-ARG achieved an F1-score of 0.795. And for the mobility identification task, we compared our PPLM with HMD-ARG and PPLM achieved an F1-score of 0.99 whereas HMD-ARG achieved a precision of 0.964 and an F1-score of 0.926.

The results of this study demonstrate several key points. First, the superior performance of PPLM across multiple tasks underscores its robustness and versatility as a pre-trained protein language model. Second, the high F1 scores indicate that PPLM is capable of maintaining a balance between precision and recall, making it highly reliable for practical applications. The consistent outperformance of PPLM across these tasks highlights its potential as a powerful tool in antimicrobial resistance research.

### Independent Test Set Validation

ProtAlign-ARG, trained on GraphPart-based clustering with a 40% similarity threshold on the HMD-ARG-DB dataset, successfully identified all novel 76 beta-lactamase genes that have been experimentally-validated by Berglund et al.^49^. Furthermore, the ARG class classification model correctly categorized these genes as beta-lactamases. To ensure that the training set did not contain these beta-lactamase genes, we conducted DIAMOND alignments, excluding any sequences with similarity above 40% during the model’s training. This performance underscores the effectiveness of our model in identifying and annotating novel ARGs compared to existing tools.

## Conclusion

Our study presents a robust pipeline and benchmarking standard for the identification and classification of ARGs and thus can contribute to advancing goals of antibiotic resistance monitoring across One Health sectors. ProtAlign-ARG introduced a novel method for taking advantage of both alignment-scoring-based tools and deep learning predictions and surpassed state-of-the-art tools in both ARG identification and class classification. One limitation of the current model is that accuracy is lower for sequences with low identity to the training data. To address this, we plan to incorporate protein 3D structure information into the models as protein structures are more conserved than sequences, potentially enhancing performance.

## Data Availability

The datasets generated during and/or analyzed during the current study are available in the ProtAlign-ARG repository - https://github.com/Shafayat115/ProtAlign-ARG.

## Acknowledgements

This work was partly funded by the National Science Foundation (NSF grants #2319522, #2125798, and #2004751).

## Author contributions statement

S.A. conducted the dataset preparation, processing, and experimentations. M.E. and N.M. assisted with the code and preprocessing of the dataset. The rest of the authors provided continuous support through suggestions and ideas and reviewed the paper.

## Additional information

**Competing interests** The authors declare no competing interests.

